# Enhanced Waddington Landscape Model with Cell-Cell Communication Can Explain Molecular Mechanisms of Self-Organization

**DOI:** 10.1101/241604

**Authors:** H. Fooladi, P. Moradi, A. Sharifi-Zarchi, B. H. Khalaj

## Abstract

The molecular mechanisms of self-organization that orchestrate embryonic cells to create astonishing patterns have been among major questions of developmental biology. It is recently shown that embryonic stem cells (ESCs), when cultured on particular micropatterns, can self-organize and mimic early steps of pre-implantation embryogenesis. A systems-biology model to address this observation from a dynamical systems perspective is essential. Here, we propose a multicellular mathematical model for pattern formation during in vitro gastrulation of human ESCs. This model enhances the basic principles of Waddington epigenetic landscape with cell-cell communication, in order to enable pattern and tissue formation. To prevent overfitting of the model, there is a minimal number of parameters in the model, which are sufficient to address different experimental observations such as the formation of three germ layers and trophectoderm, responses to altered culture conditions and micropattern diameters, and unexpected spotted forms of the germ layers under certain conditions. This model provides a basis for in-silico modeling of self-organization.

## 1 Introduction

From longtime ago, lots of prominent scientists devoted their life to think about how biological patterns emerge and how differentiation occurs. Different cell fates that organize through space and time are formed from seemingly identical ones. One of the models for explaining pattern formation was proposed by Lewis Wolpert in 1969 which is known as positional information(PI) [1]. This model suggests a specific embryonic region diffuses some morphogen, which results in a morphogen gradient. Cell fate is determined by the concentration of the morphogen that cell receives. [1]. Positional information model does not consider morphogen interactions and consequently is incapable of producing oscillatory patterns [2]. From another point of view, Alan Turing proposed reaction-diffusion(RD) model, which is based on the interaction of two morphogens and can produce oscillatory patterns [3]. Formation of different biological patterns like animal skin pattern, as well as vasculogenesis, can be justified by the RD model [2, 4, 5, 6]. Even it is possible to combine RD and PI and create another sophisticated biological model [7].

There are numerous examples of pattern formation during early human embryo development and gastrulation is among most important of them. Lewis Wolpert has mentioned “it is not birth, marriage, or death, but gastrulation which is truly the most important time in your life” [8] and this quote reveals the significant importance of gastrulation. Gastrulation is a self-organized process; which means without any external force and just by cell-cell communication, a group of identical cells differentiates and form ordered spatial pattern. During mammalian gastrulation, epiblast cells in blastocoel which had been differentiated from inner cell mass, produce three main kinds of cells: ectoderm, mesoderm, and endoderm. These three spatially organized layers are the progenitor of almost every cell in human body [9].

It is recently shown that providing geometric confinement is enough for recapitulation of ordered sequence pattern and opened new doors for studying this process more precisely in great depth [10, 11]. In this experiment, homogenous BMP4 was used as an input signal for triggering self-organized gastrulation, and cells were confined to circular micro colonies of 250 μm to 1000 μm diameter. They investigated the effect of micro pattern sizes and different concentrations of initial BMP4 on output pattern experimentally [11]. Etoc et al. followed this procedure and tried to propose a mathematical model for this qualitative behavior [12]. As a result, they found out initial receptor relocalization and interaction of BMP4 and its inhibitor, Noggin, were responsible for ordered germ layer formation. Although their suggested mathematical model was compatible with previous experiment it did not have ability to produce oscillatory pattern which was observed later [13]. Tewary et al. showed that by increasing colony diameter and initial BMP4 concentration, it is possible to observe spotty pattern [13]. They modified Etoc et al. genetic circuit and added auto-regulation link for BMP4 which makes this model capable of producing spatial oscillation and resemble Turing model in mind.

Despite the fact that their model can reproduce all the experiments, It is difficult to biologically interpret the model parameters. Correspondence between parameters and specific biological or molecular property is not clear. This restricts us from designing a new experiment to verify the applicability of this model because we do not know how one parameter variation maps to one specific factor in the lab. Furthermore, another thing that makes this model inconvenient from our perspective is ignorance of cell identity. The previous model([13]) considers BMP4 and Noggin as a continuous field and their interaction makes the sequential ordered pattern. But a more precise and realistic model should include cells as the core element and consider cell-cell communication.

We propose a mathematical model based on dynamical system theory for justifying pattern formation during in-vitro hESC gastrulation. This model not only is compatible with all the previous experiments but also considers cells as the main elements and contains parameters which have very precise biological interpretation. It also gives us an opportunity of making testable biological statements about how the pattern changes in response to changing parameters.

## 2 Methods

We proposed a model based on dynamical systems approach. The minimal genetic circuit has been considered within each cell and this circuit contains interaction of BMP4 and Noggin genes which are among the most important genes in human gastrulation. In particular, BMP4 activates Noggin and its own production and Noggin inhibits production of BMP4. It is important to note that BMP4 does not interact with Noggin directly; it first stimulates phosphorylation of Smad1 and complex of Smad1 and Smad4 transfer to the nucleus and activate Noggin expression[14]. This circuit can be considered as a dynamical system and the underlying equations that capture these interaction can be extracted. Each gene has a degradation rate as well as production rate.

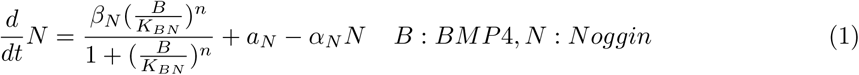

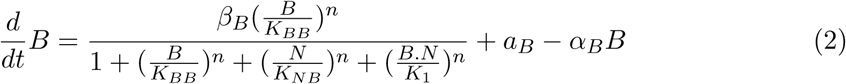

In this equation, N is the concentration of Noggin and B is the concentration of BMP4 in one cell. *α_B_* and *α_N_* stand for degradation rates of BMP4 and Noggin in each cell respectively. Furthermore, *β* and *a* can be considered as the production rates and *K* represents binding affinity of one protein to the specific promoter.

It is reasonable idea to use non-dimensional version of dynamical system equations. Nondimensionalization often reduces number of free parameters and consequently reduces search space. It is important to choose appropriate dimensionless parameters. We chose dimensionless parameters as the following:

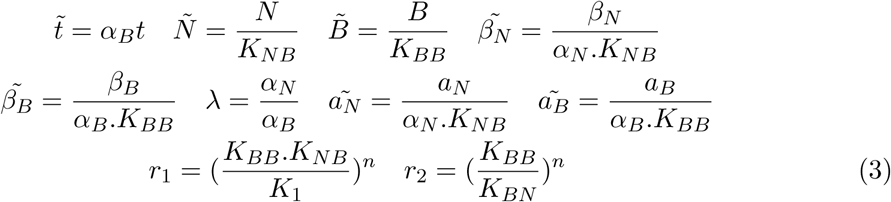

Manipulating the original equations with dimensionless parameters can lead to following result.

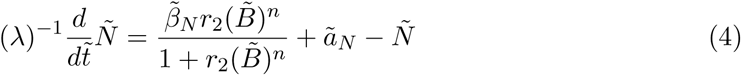

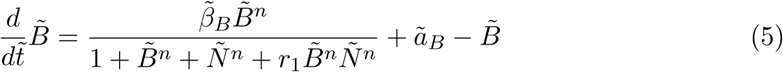

After that group of cells have been considered and within each cell this genetic circuit exists. It is possible that each protein leaves the cell and diffuses through extracellular space(Fig. 2).

## 3 Results and Discussion

In the last section, we determined the dimensionless equation of genetic circuit depicted in Fig.1. It can be seen that there are eight free parameters in this model and searching through parameters space can lead to different qualitative behaviors. To make this point clear, we simulated and tested about 100 different sets of parameters and decided to show the results for two of them (Table. 1). It is important to note that for sake of simplicity, we have omitted ~ in the following essay and all the subsequent analysis is based on the dimensionless equation.

**Table 1:** Two sets of parameters that can cause two completely different qualitative behavior.

| $a_N$ | $a_B$ | $\beta_B$ | $\beta_N$ | $\lambda$ | $r_1$ | $r_2$ | $n$ |
| --- | --- | --- | --- | --- | --- | --- | --- |
| 0.1 | 1 | 100 | 10 | 3 | 0.1 | 1 | 2 |
| 0.1 | 0.1 | 20 | 20 | 0.4 | 0.1 | 1 | 2 |

**Figure 1:**
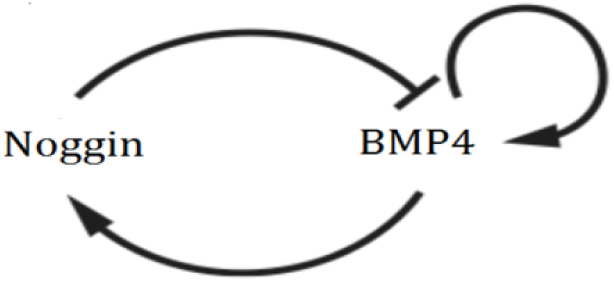
Genetic circuit that shows interaction of BMP4 and Noggin genes.

**Figure 2:**
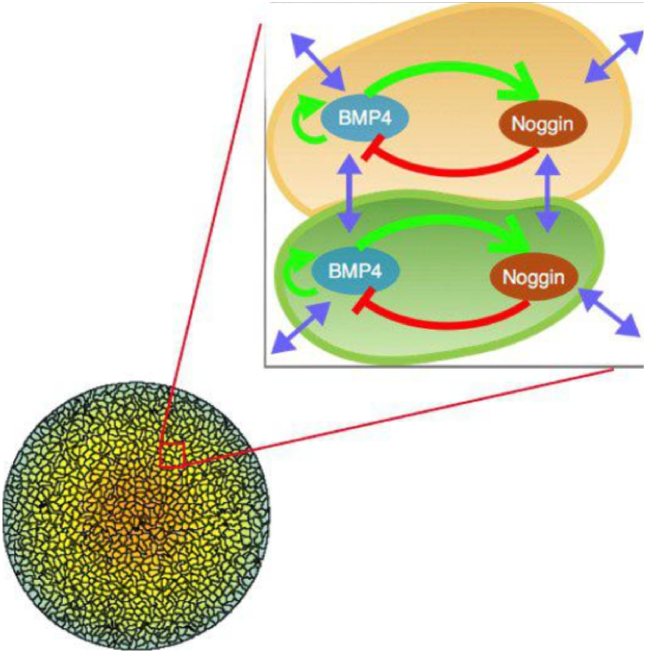
Schematic of our proposed model.

The system approaches a fixed point for the first set of parameters and it shows oscillatory behavior for the second set of parameters(Fig. 3). Even this very simple system within one cell can exhibit numerous qualitative behaviors. Our goal in this paper is considering the population of cells and placing this genetic circuit in each cell. BMP4 and Noggin proteins can leave the surface of the cells and diffuse through extracellular space and consequently influence the behavior of other cells. In this manner, we have considered cellular interaction in our model which is the key part in pattern formation.

**Figure 3:**
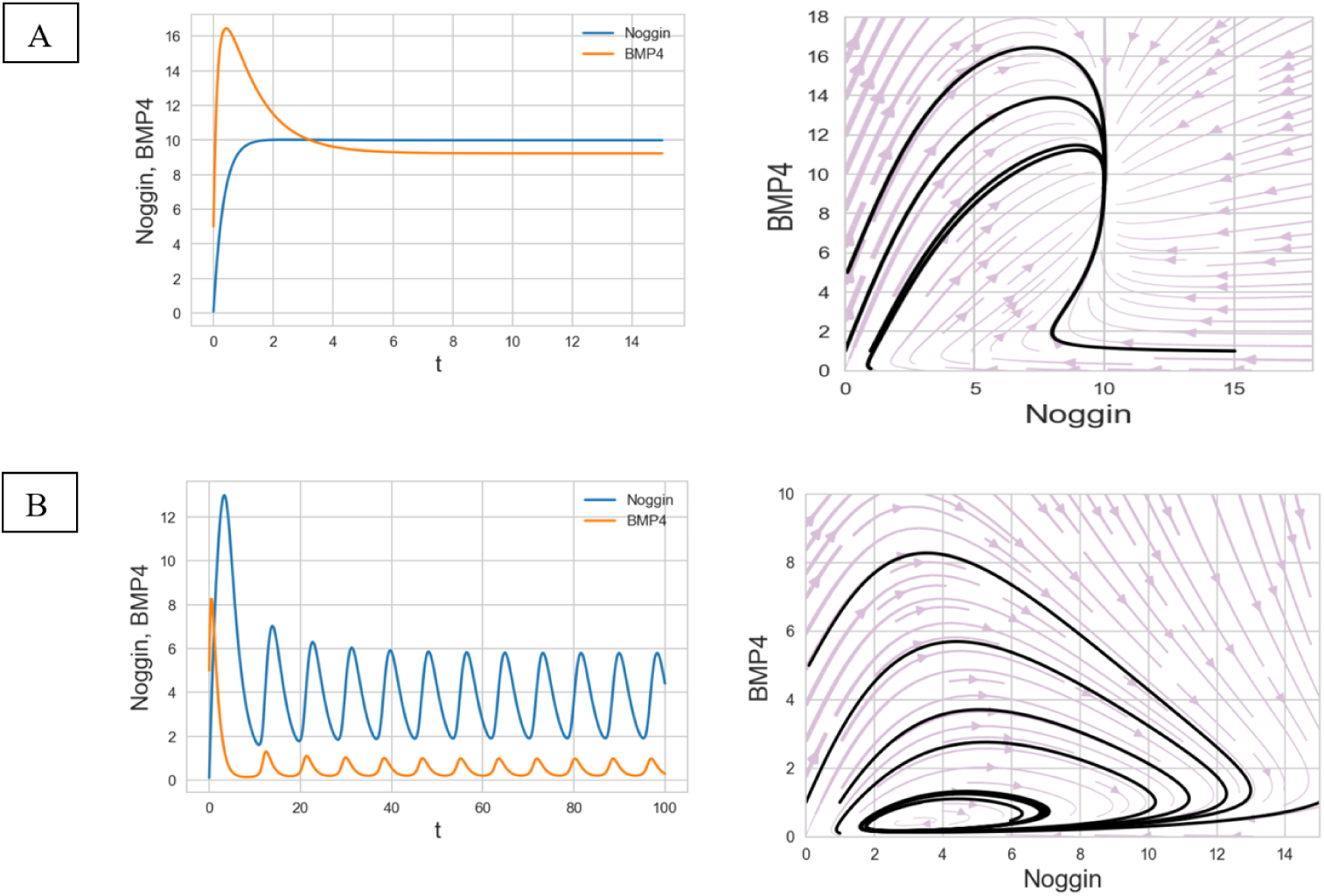
Behavior of the genetic circuit in time and state space. A) Evolution of the system when the parameters have been selected based on first row of table 1. B) Evolution of the system when the parameters have been selected based on second row of table 1. Left: Time space. Right: State space with different initial conditions.

All the simulations in the multicellular system have been done with Morpheus. Morpheus is user-friendly software designed for simulating and studying multicellular systems [15]. As we saw, our proposed model has some free parameters which must be tuned. Exhaustive search and simulating the vast number of different models, we came up with the parameters in the Table2 which cause oscillatory behavior in one cell without the presence of interaction with other cells.

Adding interaction has a significant effect on final results which can reveal the importance of interaction and diffusion to pattern formation. Furthermore, diffusion of BMP4 and Noggin is another factor that has an enormous effect on the pattern; it has been observed that desired pattern emerges when diffusion of Noggin is significantly greater than BMP4. In Fig. 4 you can observe the amount of BMP4 and Noggin within each cell and also BMP4-Noggin field in space

**Figure 4:**
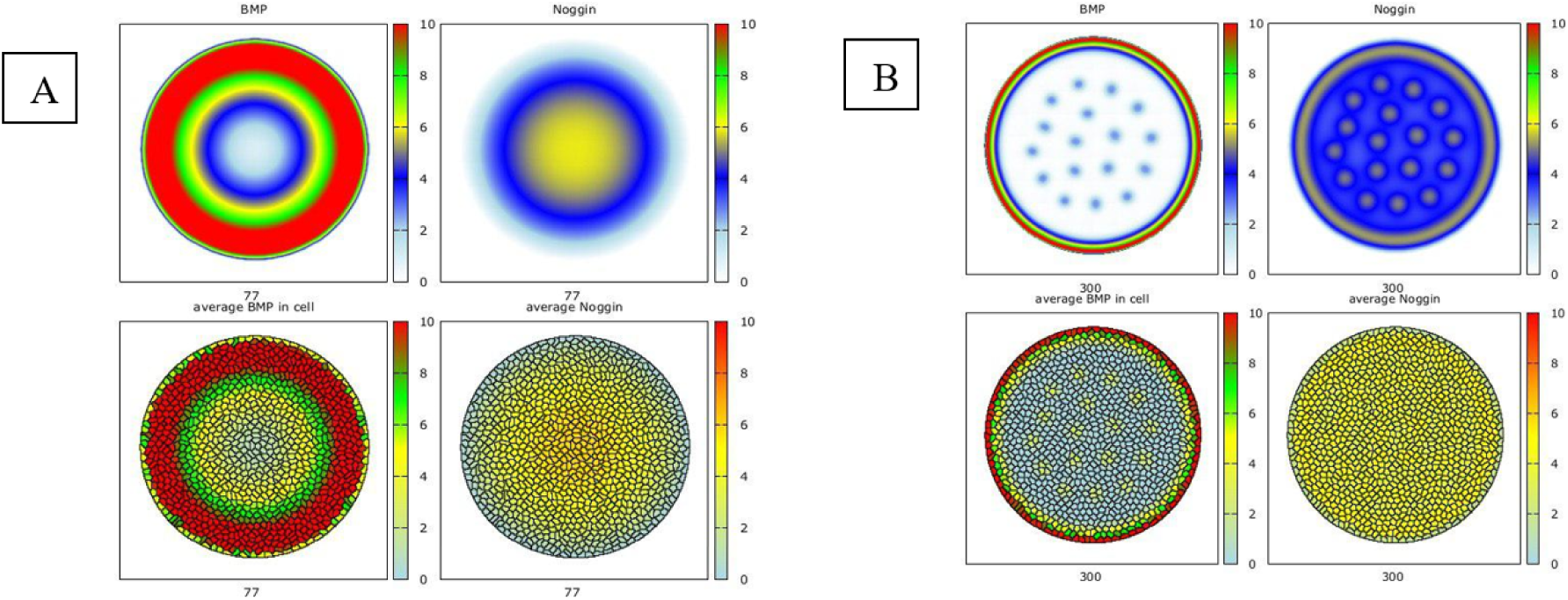
Simulation of proposed model in population of cells when the interaction and diffusion is present. These patterns have emerged after the transition time is over and when the system has reached the stable situation. A) Simulation on 500 μm colony. B) Simulation on 2.5 mm colony. Above: BMP4 and Noggin field in the environment. Below: Amount of BMP4 and Noggin within each cell.

Based on the previous experiments, we expect BMP4 to decrease from the edge towards the center of the colony, while Noggin shows the opposite behavior in 500 μm colony. After formation of BMP4-Noggin pattern, the appearance of different cell fates and ordered germ layers can be explained based on positional information mechanism.

In addition, it has been reported that by increasing colony diameter to around 3 mm, the spotty pattern will appear. In fact, our model leads to same observation and we can see the desired pattern after simulation (Fig. 4). For spotty pattern, we saw that spots appear and oscillate through time until the system reaches stable point and oscillation in time vanishes. It is important to note that all the simulations have been performed with constant boundary conditions where we assume amounts of BMP4 and Noggin in the boundaries are constant. Furthermore, it is good to mention that parameters vary within a reasonable biological range.

**Table 2:** Appropriate parameters for emergence of desired pattern.

| $a_N$ | $a_B$ | $\beta_B$ | $\beta_N$ | $\lambda$ | $r_1$ | $r_2$ | $n$ | $\tilde{D}_B = \frac{D_B}{\alpha_B}$ | $\tilde{D}_N = \frac{D_N}{\alpha_B}$ |
| --- | --- | --- | --- | --- | --- | --- | --- | --- | --- |
| 0.1 | 0.1 | 20 | 20 | 0.4 | 0.1 | 1 | 2 | 250 | 12 500 |

### 3.1 Effect of each parameters on pattern

One key advantage of our proposed model is providing a biological interpretation for each parameter. In other words, each parameter corresponds to the specific biological process or molecular properties. For example λ is the ratio of degradation of Noggin and BMP4 and varying λ means varying degradation rate of one of these proteins. We examined the effect of each factor on the resulting pattern and found that some parameters have more importance on the formation of desired pattern and system is more sensitive to their variation. We observed that the system shows more sensitivity to *r*_1_*, λ, D_BMP4_* and *D_NOG_* and is robust to variation of other parameters such as *r*_2_. For making this point clear, you can observe the effect of diffusion on patterns in 500 μm colony in Fig. 5.

**Figure 5:**
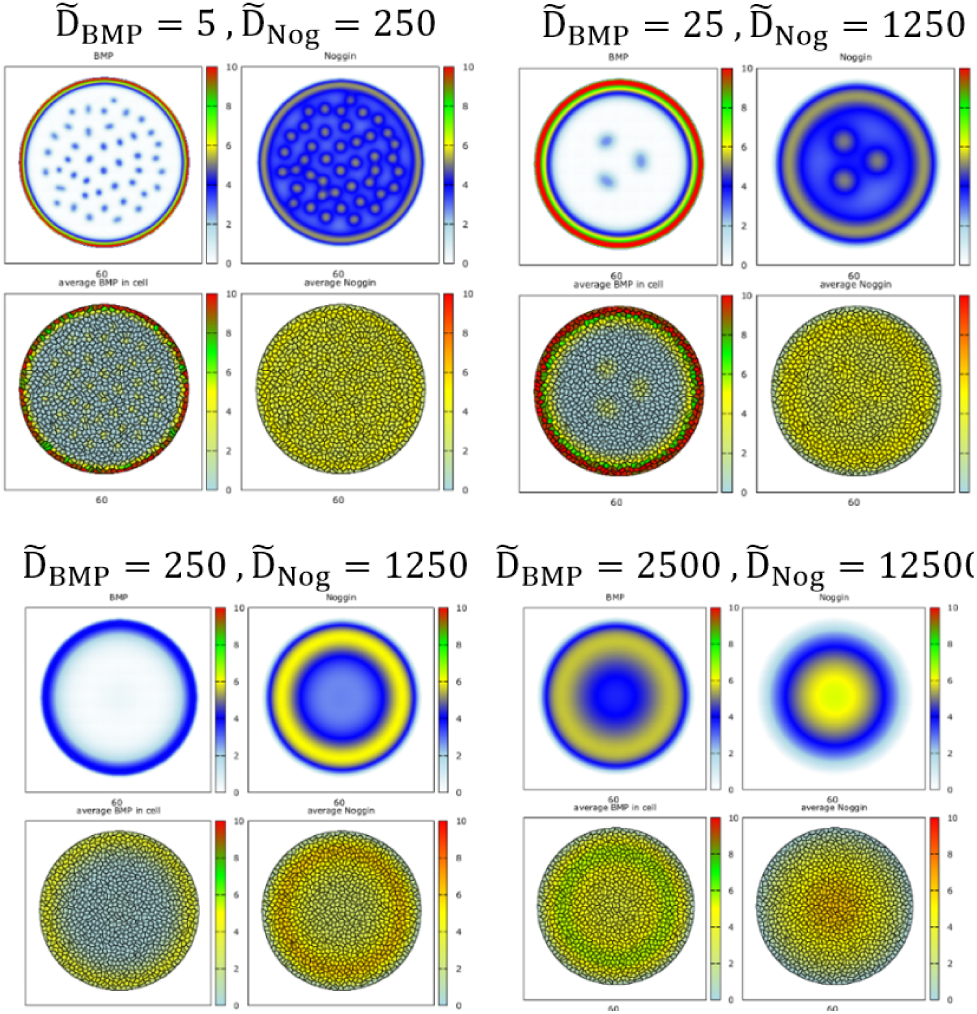
Effect of diffusion on pattern formation in 500 μm colonies. It is obvious that variation of diffusion has significant effect on the final pattern.

### 3.2 Effect of colony geometry on pattern

All the previous experiments have been done on circular confinement for colonies. We contemplated that geometry of colony should have a significant effect on the system behavior and decided to evaluate our hypothesis by simulation.

Different geometries for colonies were considered and for each one simulation performed. We decided to show the result of one geometry which resembles in vivo gastrulation. Based on simulations, our hypothesis seems reasonable; but it needs an in-vitro experiment for confirmation. We think it would be interesting to write state of the system or fate of each cell as a function of geometry or at least some features of geometry like curvature. It can be considered as future works, but for now, we have just performed some simulations on different geometries to clarify our claim.

## 4 Conclusions

We observed that cellular interaction play an important role in pattern formation; in absence of interactions, the system exhibits time-course oscillatory behaviors. In presence of such interactions, the system stabilizes. Hence, the cellular interactions are among important factors that lead the system toward the desired pattern. In addition, the colony diameter has a significant impact on the pattern that emerges; increasing colony diameter can result in the formation of the spotty pattern and our model can capture this behavior. Furthermore, we studied the effect of different parameters and observed that some of such as *r*_1_*, λ, D_B_* and *D_N_* have greater impacts on final results. Subsequently, we studied the effect of colony geometry on pattern and as we expected, even a minor change in geometry can have a significant impact on the pattern. Experimental results are required to validate our observations.

**Figure 6:**
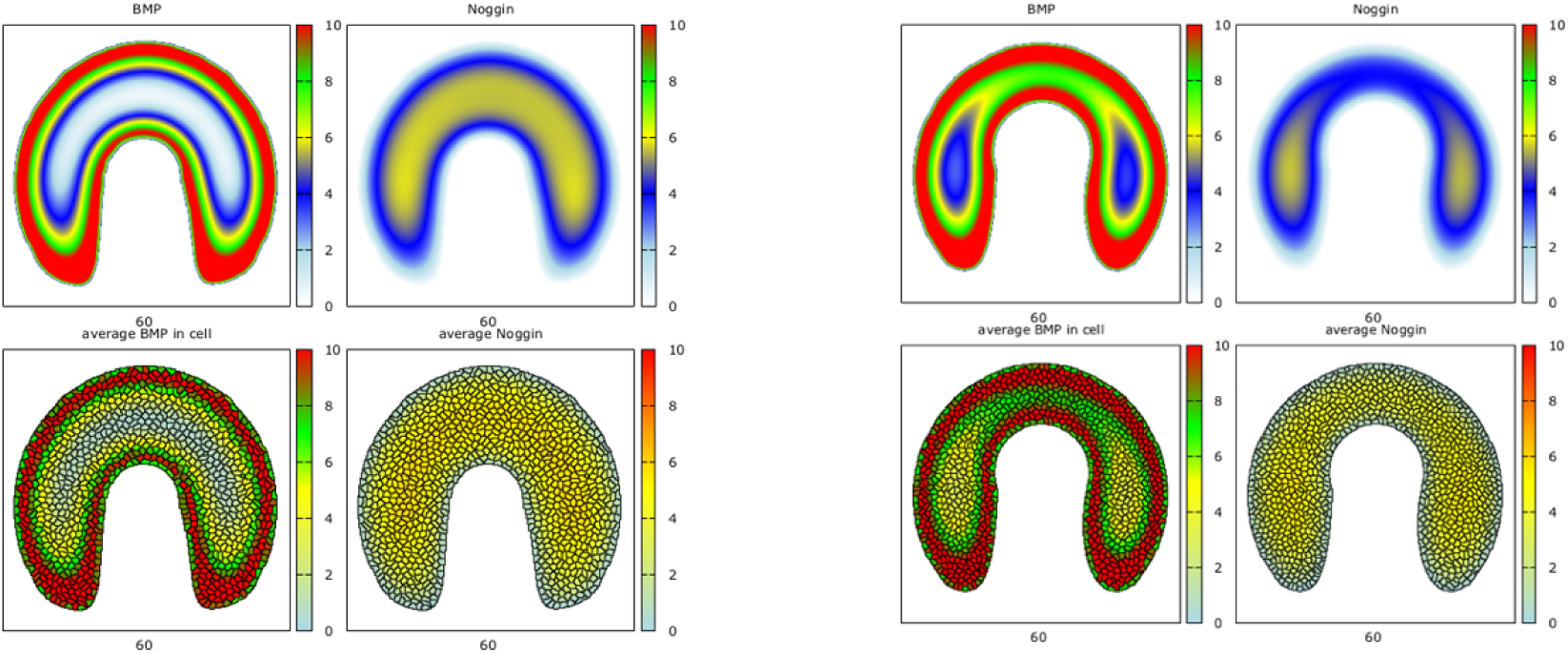
Effect of different geometric confinements on pattern formation and cell fate decision. Minor alternations of in geometry has made dramatic changes in pattern. In the right figure, the bottleneck is slightly narrower than the left figure, leading to significant change in the resulting pattern.

